# BrainBeacon: A Cross-Species Foundation Model for Single-cell Resolved Brain Spatial Transcriptomics

**DOI:** 10.1101/2025.07.08.663729

**Authors:** Chengming Zhang, Yiwen Yang, Yifeng Jiao, Qianqian Yang, Xin Guo, Jing Xu, Jiyang Li, Yuanyuan Zhou, Zhen Liu, Yushuai Wu, Yu Zhang, Chen Jiang, Jianyun Ma, Yuxuan Liu, Yao Zhang, Kai Xiao, Yidi Sun, Cirong Liu, Limei Han, Yuan Qi, Kazuyuki Aihara, Chengyu Li, Yuan Cheng, Luonan Chen, Wu Wei

**Affiliations:** Lingang Laboratory, Shanghai 200031, China; Shanghai Academy of Artificial Intelligence for Science, Shanghai 200003, China; School of Life Sciences and Biotechnology, Shanghai Jiao Tong University, Shanghai 200240, China; International Research Center for Neurointelligence, The University of Tokyo Institutes for Advanced Study, The University of Tokyo, Tokyo 113-0033, Japan; School of Mathematical Sciences and School of AI, Shanghai Jiao Tong University, Shanghai 200240, China; School of Life Science and Technology, ShanghaiTech University, Shanghai 201210, China; Key Laboratory of Systems Biology, Shanghai Institute of Biochemistry and Cell Biology, Center for Excellence in Molecular Cell Science, Chinese Academy of Sciences, Shanghai 200031, China; Artificial Intelligence Innovation and Incubation Institute, Fudan University, Shanghai 200433, China; Zhongshan Hospital, Fudan University, Shanghai 200433, China; Key Laboratory of Systems Health Science of Zhejiang Province, School of Life Science, Hangzhou Institute for Advanced Study, University of Chinese Academy of Sciences, Chinese Academy of Sciences, Hangzhou 310024, China; CAS Key Laboratory of Computational Biology, Shanghai Institute of Nutrition and Health, University of Chinese Academy of Sciences, Chinese Academy of Sciences, Shanghai 200031, China; Institute of Neuroscience, State Key Laboratory of Genetic Evolution & Animal Models, Center for Excellence in Brain Science and Intelligence Technology, Chinese Academy of Sciences, Shanghai, 200031, China

## Abstract

The brain’s functional complexity emerges from spatiotemporal heterogeneity in molecular, cellular, and circuit organization, shaped by evolution, aging, and disease. Single-cell spatial transcriptomics provides a transformative framework to decode this complexity, resolving cell-type-specific expression patterns, region-specific regulation, evolutionary divergence, and disease-associated disruptions. Despite recent advances in spatial transcriptomics, a unified foundation model capturing cross-species, whole-brain cellular architecture remains lacking. Here, we introduce BrainBeacon, the first cross-species foundation model for whole-brain spatial transcriptomics. Trained on 133 million spatially resolved cells covering a total area of 210194 mm^2^ from human, macaque, marmoset and mouse brains profiled across five major platforms, BrainBeacon integrates gene expression rankings, spatial cellular organization, and evolutionarily conserved genetic relationships into a unified transformer-based architecture. A two-stage training strategy captures both intra-cellular transcriptional dependencies and inter-cellular spatial interactions, yielding biologically grounded and interpretable representations of brain structure. Subsequently, BrainBeacon enables cross-species alignment of brain cell types and anatomical regions, facilitating integrative analyses of conserved and divergent cellular architectures. Moreover, it uncovers spatial regulatory mechanisms of aging by predicting bidirectional perturbation responses between cells and their local niche. Together, BrainBeacon serves as a foundational cross-species model for whole-brain spatial transcriptomics, enabling virtual brain atlas construction, systematic in silico perturbation studies, and mechanistic insights into brain disease pathogenesis.

## Introduction

The mammalian brain’s remarkable complexity is encoded in the spatial arrangement and molecular identity of billions of cells. A central goal of modern neuroscience is to construct comprehensive cellular atlases that link this architecture to brain function, evolution, and disease. Recent, large-scale efforts by consortia such as the BRAIN Initiative Cell Census Network (BICCN) and others have generated unprecedented transcriptome datasets of millions of individual cells across numerous brain regions in key species, including the mouse^1^, marmoset^2^, macaque^3^, and human^4^. These atlases are beginning to reveal the conserved and divergent principles of brain organization and provide a foundation for understanding the cellular basis of neurological and psychiatric disorders. However, the sheer scale and heterogeneity of these data spans multiple species, molecular modalities, and high-resolution spatial technologies (e.g., MERFISH^5^, STARmap^6^, Xenium^7^, Slide-seqV2^8^ and Stereo-seq^9^), presenting a formidable integration challenge that current computational approaches have yet to fully resolve.

The rapid progress in large-scale knowledge-infused representation learning has enables foundation models to learn fundamental principles from massive, unlabeled datasets^10^. This success highlights a critical opportunity and reveals a corresponding gap in neuroscience, as preliminary models have been developed for single-cell transcriptomics, such as Geneformer^11^ and scGPT^12^, which are trained exclusively on human data and lack spatial context. UCE^13^ extends single-cell modeling to cross-species settings, but does not incorporate spatial information. GeneCompass^14^ leverages large-scale single-cell transcriptomes from human and mouse to model cross-species gene regulatory mechanisms using prior biological knowledge, but does not incorporate spatial information. In contrast, Nicheformer^15^ jointly models spatial and single-cell transcriptomics across the same species. CellPLM^16^ integrates spatial data but does not explicitly model cross-species variation. No existing framework jointly captures the structured spatial architecture and the deep evolutionary conservation spanning rodents and primates across the whole brain, underscoring the need for a generalizable and interpretable model tailored to spatial neuroscience.

To address this critical gap, we present BrainBeacon, the first brain-specific foundation model designed to unify cross-species and cross-platform spatial transcriptomics data into a single, coherent framework. Beyond modeling gene expression ranking, BrainBeacon encodes each cell with a composite token that integrates spatial neighborhood-level expression deviations, local cell density, and cross-species homology features, thereby enabling cross species generalization in spatial transcriptomic contexts. We pretrain BrainBeacon on a comprehensive corpus of 92,076 gene vocabulary from 133 million cells, encompassing human, mouse, macaque, and marmoset brain samples collected using five major spatial platforms: MERFISH^5^, STARmap^6^, Xenium^7^, Slide-seqV2^8^ and Stereo-seq^9^. BrainBeacon employs a two-stage training strategy that first models intra-cellular gene dependencies through masked gene modeling, and subsequently incorporates inter-cellular spatial context to capture local microenvironmental structure. This dual design facilitates both robust representation learning and explicit spatial reasoning across the most heterogeneous brain datasets available to date.

We demonstrate that BrainBeacon exhibits powerful generalization across a wide range of downstream tasks, establishing a new standard for computational neuroanatomy. The model enables high-accuracy spatial cell clustering across species and platforms, outperforming existing methods in challenging transfer-learning settings such as snRNA-seq-to-spatial integration and cross-species label transfer. By analyzing its learned representations, BrainBeacon uncovers conserved and species-specific cell types and spatial architectures, elucidating fundamental evolutionary principles of primate and rodent brain organization. Finally, we introduce a novel in silico perturbation framework that leverages the model’s predictive power to simulate the effects of both cell-intrinsic and microenvironment-level interventions on gene expression in a spatial context. Together, these results establish BrainBeacon as a unified, extensible, and interpretable foundation model for decoding the conserved and dynamic principles of brain organization.

## Results

### A foundation model for brain spatial transcriptomics

To capture fine-grained spatial transcriptomic patterns across species and platforms, we developed BrainBeacon, a dual-stage transformer framework trained on a comprehensive spatial brain atlas (BrainST-133M), comprising 133 million cells, covering a total area of 210194 mm^2^, from four species and five spatial profiling technologies **(Fig. 1a**). Each cell is represented by multimodal features, including gene expression, spatial coordinates, local cell density, gene deviation relative to its local niche, and metadata such as species and platform identity (**Fig. 1b**). BrainBeacon consists of two sequential modules: an intra-cell transformer, which models gene-level structure and predicts masked gene tokens based on identity, evolutionary features (e.g., ESM2 sequence embedding, homology group), and RNA type; and an inter-cell transformer, which models spatial neighborhoods within each slice to reconstruct cell-level expression profiles (**Fig. 1c**). The output of the second stage corresponds to predicted gene expression counts modelled using a negative binomial distribution, capturing the overdispersed nature of transcriptomic data. This architecture enables integrated representation learning across spatial, species, and molecular dimensions, supporting diverse downstream tasks such as spatial cell clustering, cross-species label transfer, and spatial in-silico perturbation inference (**Fig. 1d-f**).

**Fig. 1:**
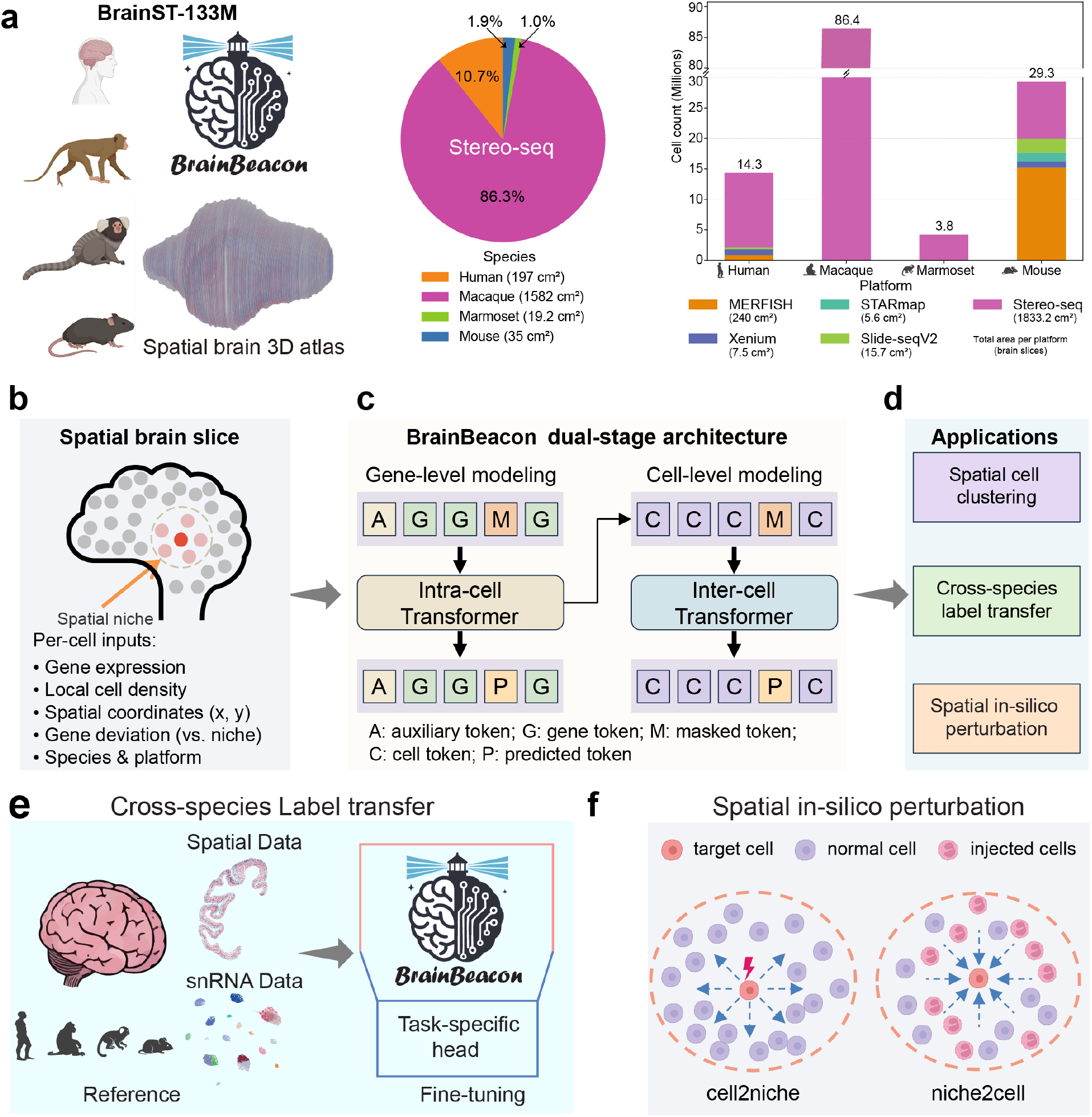
Overview of the BrainBeacon framework and architecture. **a**, Spatial brain atlas dataset (BrainST-133M) comprising 133 million cells across four species (human, macaque, marmoset, mouse) and five spatial transcriptomics platforms (MERFISH, Xenium, STARmap, Slide-seqV2, Stereo-seq). Left, pie chart shows the proportion of Stereo-seq–acquired brain slice area (in cm^2^) across species.Right, bar plot summarizes cell counts and total imaged brain slice area per platform (in cm^2^), highlighting the dominance of Stereo-seq in both cell yield and coverage. **b**, Each cell is represented by multiple features, including gene expression, spatial coordinates, local cell density, gene deviation (relative to surrounding niche), and species/platform identity. **c**, BrainBeacon adopts a dual-stage transformer architecture: intra-cell modeling at the gene level and inter-cell modeling at the cell level. Intra-cell modeling predicts masked gene tokens within a cell; inter-cell modeling predicts masked cell expression within a spatial slice. **d**, BrainBeacon supports diverse downstream tasks, including spatial cell clustering, cross-species label transfer, and spatial in-silico perturbation. **e**, Cross-species label transfer is achieved by fine-tuning BrainBeacon on spatial or snRNA reference data and applying it to spatial data across species. **f**, Spatial in-silico perturbation strategies modeled by BrainBeacon. In“cell2niche”, specific cells are perturbed and effects on surrounding niche are observed. In “niche2cell”, local microenvironments are perturbed to assess their influence on central target cells.

### BrainBeacon enables accurate discrimination of brain cell types and spatial domains

To evaluate the zero-shot representation capabilities of BrainBeacon on spatial transcriptomic data, we systematically benchmark the model across multiple publicly annotated datasets covering all five platforms and four species. BrainBeacon consistently outperforms competing methods across all datasets, with adjusted Rand index (ARI) values reaching up to 0.79 at the subclass level. CellPLM also demonstrates strong performance, maintaining high clustering accuracy with a maximum ARI of 0.75. Owing to its dual-level modeling of both cells and genes, BrainBeacon further surpasses other models in silhouette score (ASW), reflecting superior cluster compactness and separation (**Fig. 2a**).

**Fig. 2:**
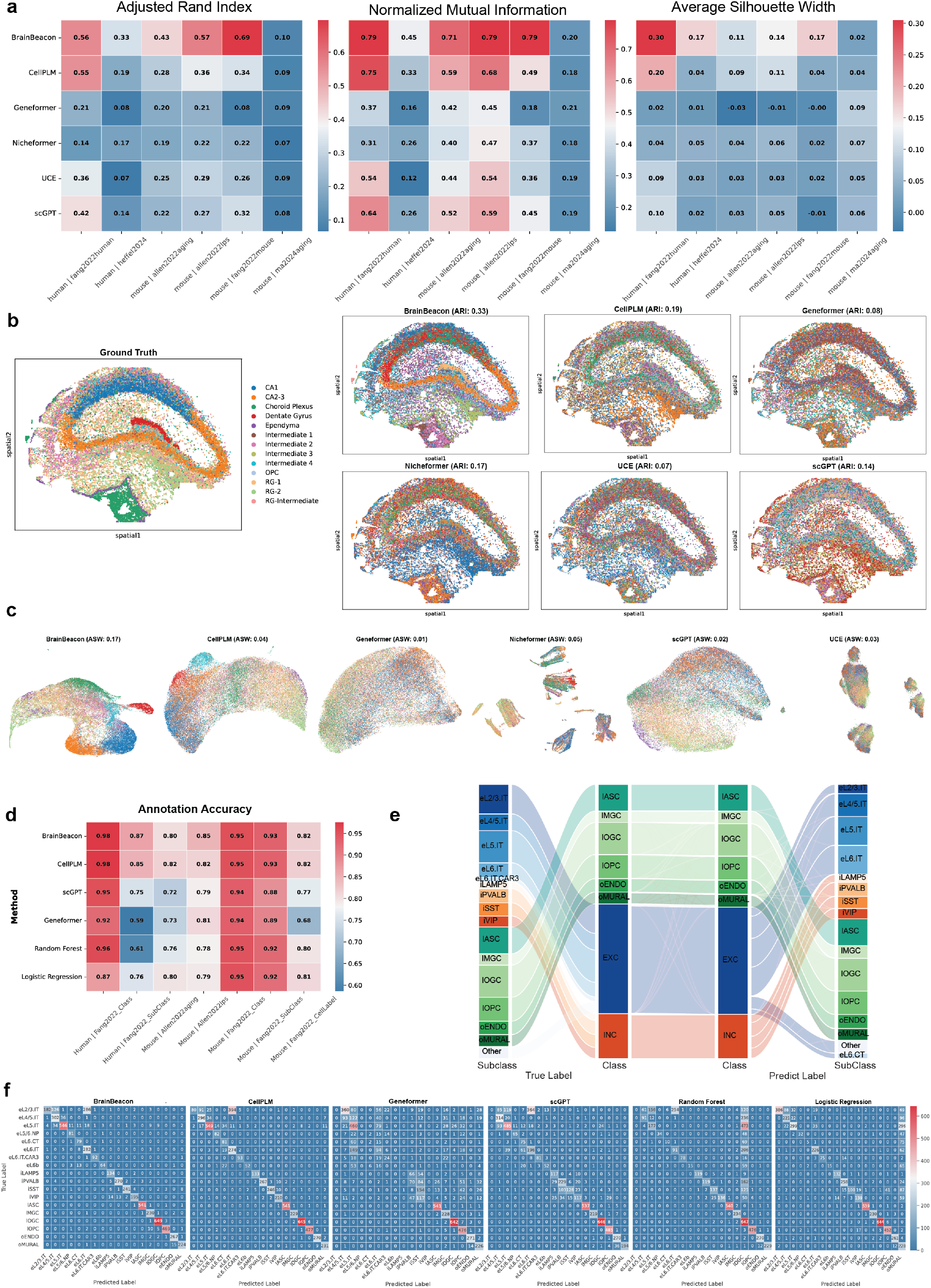
Zero-shot representation and annotation performance of BrainBeacon across spatial transcriptomics datasets. **a**, Quantitative evaluation of unsupervised clustering performance on six benchmark datasets across three metrics: adjusted Rand index (ARI), normalized mutual information (NMI), and average silhouette width (ASW). BrainBeacon consistently achieves the highest scores across datasets and metrics, especially at the subclass level. **b**, Spatial visualization of hippocampal tissue from the Ma2024Aging dataset. BrainBeacon embeddings reconstruct expected laminar organization of the CA1, CA2-3, and Dentate Gyrus regions. **c**, Uniform Manifold Approximation and Projection (UMAP) plots showing the learned embedding space from five representative datasets. BrainBeacon representations exhibit clear separation of major neuronal and glial populations, reflecting biologically meaningful structures. **d**, Cell type annotation accuracy in zero-shot label transfer across tissue slices. BrainBeacon achieves the highest accuracy at both class and subclass levels, maintaining robust performance even with increased subtype granularity. **e**, Sankey plot showing subclass-level label correspondence between ground truth and BrainBeacon predictions. Most subclasses are accurately assigned; misclassifications mainly occur between biologically similar types such as iVIP and iLAMP5. **f**, Confusion matrices for subclass predictions across five hippocampal slices. BrainBeacon achieves diagonally dominant matrices, indicating high precision; most errors are restricted to closely related subtypes (e.g., eL4/5.IT vs. eL5.IT).

In situ spatial distributions show that BrainBeacon accurately identifies key hippocampal regions, such as the CA fields and the Dentate Gyrus, with strong agreement between predicted cell types and anatomical structures (**Fig. 2b**). These spatial patterns suggest that BrainBeacon captures biologically meaningful regional identities, consistent with known hippocampal organization.

To further examine cell-type resolution in the latent space, we visualize UMAP projections of cell embeddings from BrainBeacon and other baseline models (**Fig. 2c**). BrainBeacon’s embedding space exhibits clear separation of major neuronal subtypes, including well-defined and spatially coherent clusters for CA1, CA2-3, and Dentate Gyrus neurons, indicating its ability to distinguish mature excitatory lineages. Notably, the radial glia subtype RG-2 also forms a distinct, isolated cluster with minimal overlap with other glial types, reflecting its unique transcriptional profile. These results indicate that BrainBeacon’s latent representation not only preserves developmental lineage continuity but also enhances subtype-level resolution across heterogeneous brain cell populations.

In comparison, CellPLM preserves a smoother global structure but lacks fine-grained separation of cell types, failing to resolve key subpopulations such as CA neurons and RG variants. Geneformer exhibits blurred cluster boundaries with limited discriminability. Both

Nicheformer and UCE over-segment the data, producing artificially sharp clusters that disrupt lineage relationships. scGPT performs moderately, resolving a few distinct cell types (e.g., CA1, Ependyma), but struggles to separate others, particularly across the CA-DG axis and within glial subtypes. Collectively, BrainBeacon achieves a better balance between cell-type resolution and lineage continuity, outperforming other models in accurately representing both local subtype structures and global tissue context.

To further evaluate model performance on the task of spatial cell type annotation, we test its ability to transfer labels across slices. Specifically, we select publicly available datasets with high-quality expert annotations spanning multiple tissue sections. In each case, one annotated slice serves as the reference, and labels are transferred to other slices in a zero-shot fashion. Under this realistic setting, BrainBeacon achieves remarkably high accuracy at the class level, exceeding 98% on average. Even in challenging scenarios involving nearly 30 subclasses, it maintains an accuracy of ~87%, demonstrating strong generalization and fine-grained discriminative power. Notably, CellPLM—leveraging neighborhood-aware modeling—also yields competitive results, while both models significantly outperform Geneformer, scGPT, and classical machine learning baselines (**Fig. 2d**).

The Sankey plot (**Fig. 2e**) shows that BrainBeacon achieves near-perfect classification for major cell classes, including Excitatory (EXC), Inhibitory (INC), and Non-Neuronal populations, with very few misclassifications. At the subclass level, the model maintains strong overall performance; however, certain types such as iLAMP5 and iVIP exhibit moderate misassignments, which may stem from overlapping gene expression profiles or diffuse spatial localization. In comparison, subtypes such as eL2/3.IT, iSST, and oMURAL are classified with high accuracy and consistency across anatomical sections.

Confusion matrix analysis (**Fig. 2f**) further reveals that BrainBeacon’s predictions are strongly concentrated along the diagonal, indicating precise subclass-level correspondence. Most misclassifications occur between closely related lineages (e.g., eL5.IT vs. eL4/5.IT, iSST vs. iVIP), aligning with biological expectations. By comparison, baseline models such as Random Forest and Logistic Regression exhibit widespread errors across subclasses, with notable category mixing. Collectively, these results demonstrate that BrainBeacon not only leads in overall annotation accuracy but also delivers superior subclass-level discrimination and robust generalization in complex spatial transcriptomics settings.

### BrainBeacon annotates brain regions through cross-species transfer learning

The advent of large-scale biological foundation models has introduced a new paradigm for addressing complex problems in life sciences. In this study, we demonstrate how a pre-trained brain foundation model, BrainBeacon, can be adapted through task-specific fine-tuning to tackle the core challenge of cross-species brain cell atlas comparison^17^. The centrepiece of our workflow, illustrated in **Fig. 3a**, is the “Reference-Guided Fine-Tuning” step. The objective of this step is to specialize the general biological knowledge contained within the BrainBeacon model by adapting it to a high-quality, spatially-resolved reference system. To this end, we constructed a high-resolution spatial cell atlas of the macaque neocortex to serve as this “teacher” or “anchor” dataset (**Fig. 3e**).

**Fig. 3:**
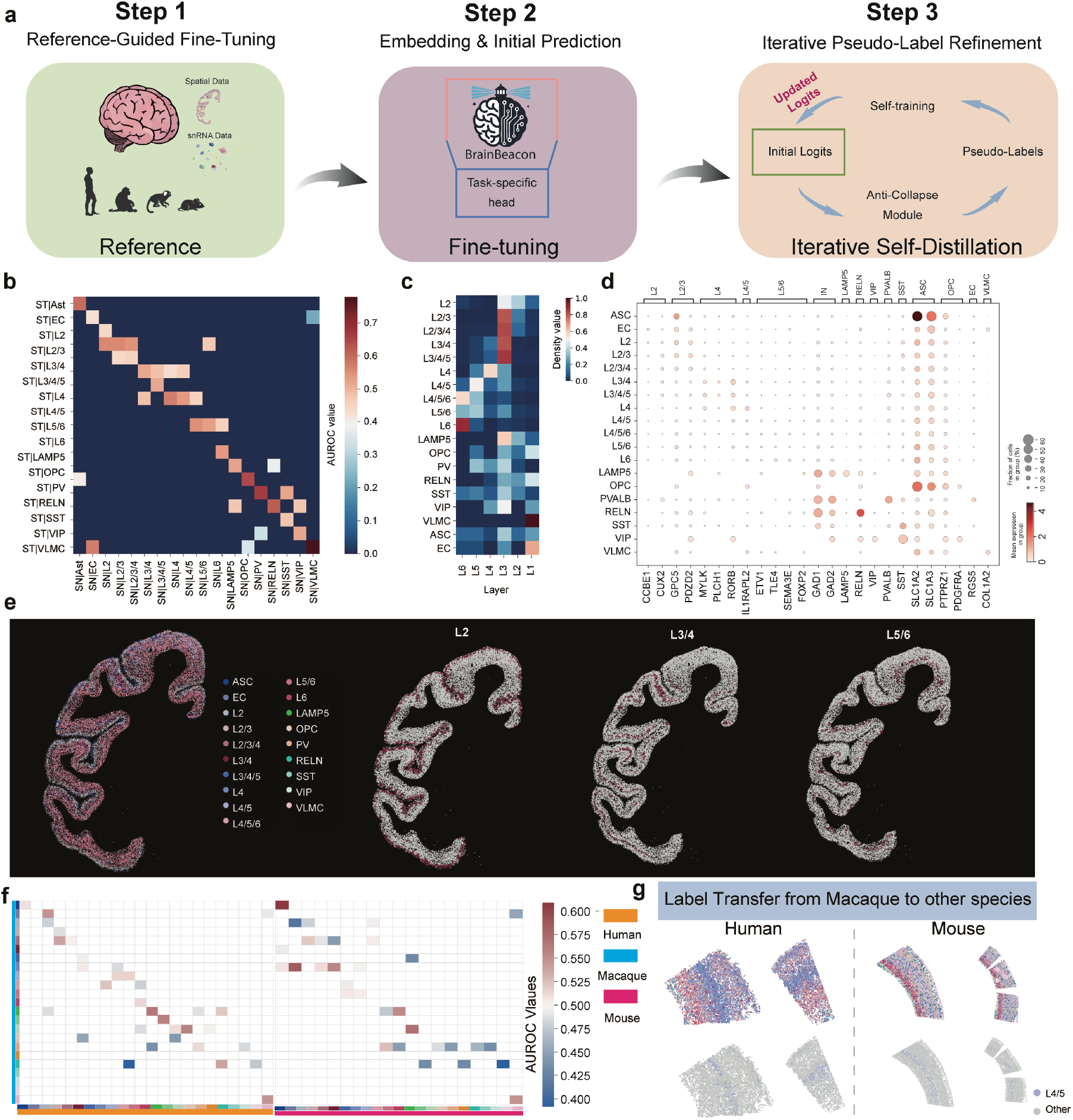
Model-driven cross-species annotation and spatial mapping of mammalian brain cell types. **a**, Schematic of the three-step computational pipeline. The model is first fine-tuned using a spatially-resolved transcriptomics (ST) reference (Step 1), followed by initial prediction on a target dataset (Step 2). Step 3 involves an iterative pseudo-label refinement process, which incorporates self-training and an anti-collapse strategy to enhance prediction accuracy. **b-d**, Intra-species validation of the model’s performance on the macaque reference. b, Confusion matrix showing high accuracy in projecting cell type labels from a macaque snRNA-seq dataset onto the macaque ST dataset. The strong diagonal indicates correct classification across excitatory neurons (e.g., L2/3, L4/5), inhibitory neurons (e.g., LAMP5, PV), and non-neuronal cells. **c**, Heatmap of laminar density, confirming that predicted excitatory neuron subclasses (L2, L3/4, L5/6) localize to their expected cortical layers. **d**, Dot plot validating predicted cell types against the expression of canonical marker genes. **e**, Spatial visualization of the annotated macaque ST reference dataset, showing the ground-truth locations of major cell types used for model fine-tuning. **f**, Cross-species performance assessment. The heatmap displays AUROC scores evaluating the model’s ability to transfer labels from macaque to human and mouse, quantifying the conservation of cell type signatures. **g**, Application of the model for cross-species annotation. The macaque-trained model successfully predicts the spatial locations of homologous cell types in human and mouse ST datasets. Lower panels highlight key examples, such as L4/5 neurons in human and mouse, demonstrating the model’s practical utility.

To validate the efficacy of our fine-tuning strategy, we first assessed the performance of the fine-tuned BrainBeacon model within the macaque species. The results indicate that the fine-tuning was exceptionally successful. The model not only projected cell type labels from single-nucleus RNA-seq (snRNA-seq) to their correct locations in spatial transcriptomics (ST) data with high fidelity (**Fig. 3b**), but its predictions were also biologically sound: the various neuronal subtypes formed a cortical laminar organization consistent with classic neuroanatomical principles (**Fig. 3c**), and their distributions showed strong concordance with their respective canonical marker gene expression patterns^18^(**Fig. 3d**). This demonstrates that our strategy successfully integrated the model’s general knowledge with the specific details of the reference atlas.

Having established the reliability of the fine-tuned model, we then unleashed its full potential to solve the critical problem of cross-species annotation. We used this macaque-tuned model to directly annotate brain ST data from human and mouse. We systematically quantified the performance of this process using AUROC scores (**Fig. 3f**). The results revealed remarkable cross-species transfer capabilities. This superior performance stems from the synergy between the general, conserved biological features pre-learned by the foundation model and the precise “anchor” provided by the high-quality reference atlas, enabling the model to “see through” superficial inter-species differences in gene expression and grasp the core essence of cellular identity.

Finally, we showcase the practical application of this powerful capability in Fig. 3g. The fine-tuned BrainBeacon model successfully constructs de novo spatial cell maps in human and mouse tissue sections, accurately identifying broad cell classes and resolving fine-grained homologous subtypes. These include Layer 4/5 neurons in human and mice, which are correctly mapped to their respective anatomical domains.

In conclusion, our work shifts the paradigm from “building one model for one task” to a more efficient approach of “adapting one foundation model for many tasks.” These findings establish that, through targeted fine-tuning on high-quality reference data, BrainBeacon can be rapidly transformed into a powerful tool for solving specific domain challenges. This paves the way for the future rapid and cost-effective construction of multi-species, multi-regional brain cell atlases and facilitates deep, comparative neuroscience research.

### Virtual spatial perturbation reveals context-aware remodeling of aging-associated transcriptional states

To investigate the causal impact of gene expression on cellular and microenvironmental states, we develop a virtual spatial perturbation framework termed cell2niche perturbation (**Fig. 4a**). This method simulates targeted gene knockouts in specific cells and evaluates the resulting changes in both the perturbed cells and their spatial neighbors within the embedding space. We select corresponding fields of view (FOVs) from young (2-month-old) and aged (25-month-old) mouse hippocampal slices as the perturbation targets (**Fig. 4b**), focusing on a curated set of aging-associated differentially expressed genes (DEGs)^19^.

**Fig. 4.**
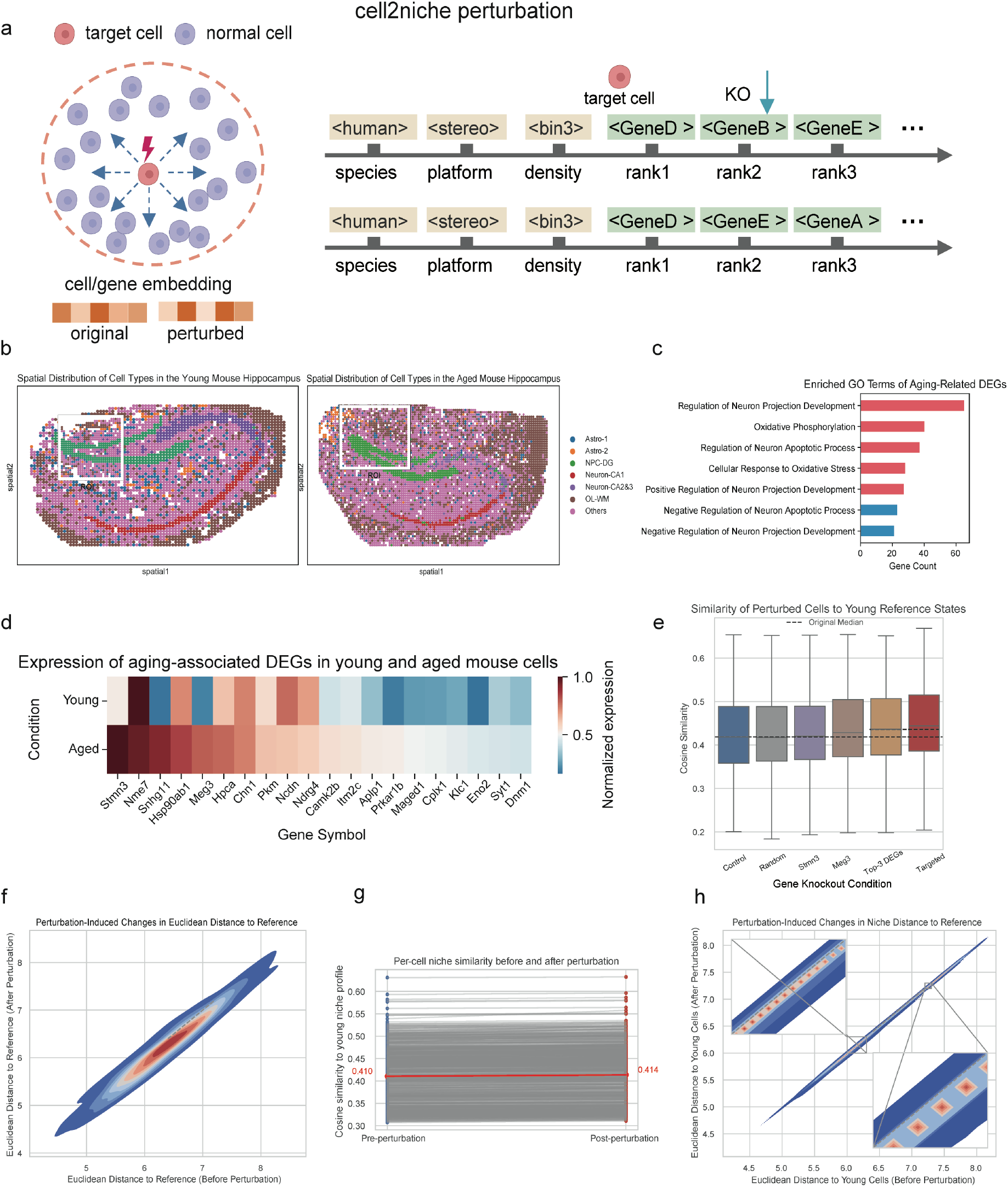
Modeling spatial gene perturbations with the cell2niche framework. **a**, Schematic of the cell2niche perturbation framework. Target cells within a spatial microenvironment are selected for in silico gene knockout by replacing ranked gene tokens in the input sequence. Perturbations are applied only to target cells (red), while spatial neighbors (blue) remain unchanged. **b**, Spatial distributions of cell types in young (left) and aged (right) mouse hippocampus, highlighting the selected FOVs used for perturbation experiments. **c**, GO enrichment of aging-related differentially expressed genes (DEGs) reveals involvement in neuronal development, apoptosis, and oxidative stress regulation. **d**, Normalized expression of aging-related DEGs in young and aged hippocampal cells. **e**, Cosine similarity between perturbed aged cells and the young reference set across different gene knockout conditions. The Meg3-Snhg11-Hsp90ab1 combination yields the highest similarity. **f**, Perturbation-induced changes in Euclidean distance to young reference cells. Triple knockout results in the largest shift toward the young state. **g**, Niche-level embedding shifts. Each line represents a target cell’s spatial neighbor; y-axis shows cosine similarity to young reference cells before and after perturbation. **h**, Euclidean distance shifts for spatial neighbors after perturbation, showing consistent neighborhood-level responses despite unmodified expression.

Gene Ontology (GO) enrichment analysis reveals that these DEGs are involved in neuronal projection regulation, apoptotic signaling, and oxidative stress responses, highlighting a global shift in pathways related to neuronal plasticity, cell survival, and mitochondrial metabolism during aging (**Fig. 4c**). These genes show consistent differential expression across young and aged hippocampal cells, including Stmn3, Meg3, Snhg11, and Hsp90ab1 (**Fig. 4d**).

We first assess the effects of single-gene knockouts. While knockout of highly expressed genes like Stmn3 resulted in minimal representational changes, Meg3 knockout significantly increased the cosine similarity between aged and young cells (**Fig. 4e**), suggesting a central role in maintaining aged-specific transcriptomic states. We next explore combinatorial perturbations and find that a triple knockout of Meg3, Snhg11, and Hsp90ab1 produce the strongest shift toward the young state, as measured by both cosine similarity (**Fig. 4e**) and Euclidean distance reduction (**Fig. 4f**). Notably, the functional relevance of this gene set spans neuronal regulation (Meg3), oxidative and proteotoxic stress response (Hsp90ab1), and inflammation-related processes (Snhg11)^20-22^.

Remarkably, although only selected cells are perturbed, the spatial neighbors that are not perturbed at the expression level also exhibit a consistent drift toward young-like states (**Fig. 4g-h**). This emergent niche-level response demonstrates that BrainBeacon captures context-aware signal propagation, enabling systematic dissection of perturbation effects across spatial tissue structures.

## Discussion

In this study, we present BrainBeacon, a cross-species foundation model for brain spatial transcriptomics that unifies gene expression, spatial context, and evolutionary priors into a scalable transformer-based architecture. Trained on over 133 million spatially resolved cells spanning four species and five platforms, BrainBeacon achieves robust performance in cell representation, cross-species cell annotation, and spatial perturbation prediction across diverse datasets.

By modeling intra-cellular gene dependencies and inter-cellular spatial interactions in a two-stage architecture, BrainBeacon captures biologically meaningful cell-cell relationships and regulatory programs. The model accurately transfers cell type labels across species, reconstructs spatially coherent regulon activity, and reveals transcriptional reprogramming associated with aging. Moreover, its virtual perturbation framework enables causal interrogation of gene function and niche-level effects in situ, offering a powerful tool for understanding the dynamics of brain aging and inflammation.

Despite these advances, BrainBeacon currently operates primarily on transcriptomic input and does not yet incorporate other modalities such as chromatin accessibility, protein expression, or histological imaging. Integrating these complementary data types will likely enhance the model’s resolution and interpretability. In particular, coupling spatial transcriptomics with co-registered histological sections may facilitate joint modeling of molecular states and tissue architecture, enabling high-fidelity reconstruction of cell–tissue interactions.

Looking forward, BrainBeacon provides a generalizable and extensible framework for decoding spatial brain biology. Its capacity to unify heterogeneous data sources across species and platforms lays a foundation for comparative neuroscience, brain atlas construction, and translational research in aging and neurodegenerative disease.

## Methods

### Datasets and preprocessing

We constructed a comprehensive single-cell spatial transcriptomic corpus of the brain to serve as the foundational dataset for training and evaluating the BrainBeacon model. This dataset encompasses samples from four species, covering human, mouse, macaque, and marmoset, and includes data generated using five spatial transcriptomics platforms: MERFISH, Xenium, STARmap, Slide-seqV2, and Stereo-seq. All datasets feature spatial coordinates and single cell resolution, enabling systematic benchmarking of the model across biological and technological contexts. Data were collected and harmonized from public repositories, published datasets, resulting in a total of 133,847,435 cells across 1,386 tissue sections from 177 donors. Notably, the macaque and mouse datasets include three and two whole-brain datasets, respectively, while the majority of remaining data correspond to cortical regions.

To ensure consistency and scientific rigor in model training and cross-task evaluation, we developed a unified preprocessing pipeline adapted to platform-specific characteristics. For high-throughput sequencing-based platforms such as Stereo-seq and Slide-seqV2, we applied stringent quality control thresholds. Specifically, we retained cells with at least 50 detected genes and excluded genes expressed in fewer than 50 cells. In addition, cells with total transcript counts below 100 were removed to eliminate low-quality or low-complexity observations. In addition, we applied neighborhood-based imputation to Stereo-seq data to enhance expression completeness. For image-based platforms, which typically provide higher per-cell expression quality such as MERFISH, STARmap, and Xenium, we applied a more relaxed quality control strategy. Specifically, we excluded cells with fewer than 20 expressed genes and removed genes expressed in fewer than 20 cells, thereby filtering out only extremely low-quality cells. To reduce platform-specific biases, we computed gene-wise expression means from healthy samples for each platform and used them to perform mean normalization independently across datasets.

#### Training corpus compilation

The final training corpus constructed for BrainBeacon comprises 85,435,334 high-quality single-cell spatial transcriptomic profiles, obtained after standardized quality control procedures. These data span 1,386 tissue sections and 177 individual donors, providing a large-scale, diverse, and high-resolution resource for model training and evaluation. Datasets were obtained from public repositories, supplementary materials. The collection spans four species, including human, mouse, macaque, and marmoset, and covers five major spatial transcriptomics platforms: MERFISH, Xenium, STARmap, Slide-seqV2, and Stereo-seq. All data were uniformly converted into the .h5ad format, preserving key modalities including expression matrices, spatial coordinates, and metadata such as platform, species, anatomical region, and donor information.

Among the species, macaque contributed the largest portion of the corpus, with 51,179,090 cells (59.90%), including an additional 27 million whole-brain cells generated using Stereo-seq. Mouse data accounted for 26,638,954 cells (31.18%), followed by human with 3,822,733 cells (4.47%), and marmoset with 3,794,557 cells (4.44%).

Platform-wise, Stereo-seq dominated the dataset with 84,386,474 cells (82.68%), primarily covering whole-brain slices from non-human primates and mice. MERFISH contributed 13,542,141 cells (13.27%) across diverse cortical regions, while Slide-seqV2, STARmap, and Xenium accounted for 1,921,166 (1.88%), 1,322,922 (1.30%), and 862,910 (0.85%) cells, respectively.

To mitigate the inherent imbalance in sample composition across species and platforms, we implemented a stratified sampling strategy during model training. Specifically, we assigned smoothed inverse-proportional sampling weights based on the number of cells within each species-platform combination, ensuring that data from underrepresented groups appeared more frequently during training. This strategy effectively reduced sampling bias, prevented overfitting to dominant platforms such as Stereo-seq or to any single species, and enhanced the model’s generalizability across cross-platform and cross-species tasks. Overall, this corpus represents the first large-scale, systematically integrated spatial transcriptomic atlas of the brain across five platforms and four species, forming a comprehensive, balanced, and structurally coherent foundation for training the BrainBeacon model.

#### Platform-specific preprocessing

Given the high sparsity of spatial transcriptomic data and the unreliability of absolute expression values at the single-cell level, we used gene expression rankings within each cell as model input. We further computed two spatially informative features: local spatial density and expression deviation relative to neighboring cells. These features capture structural variation in the tissue and are indicative of anatomical boundaries and spatially regulated gene expression patterns. These features, together with species and platform identifiers and the expression rank vectors, were used to construct the model input representation.

#### Unified gene representation and alignment

To facilitate cross-species learning, we standardized gene nomenclature by converting all gene symbols to Ensembl IDs and built a gene homology graph using orthology relationships obtained from the BioMart database. We further encoded genes using embeddings derived from the ESM2 protein language model, enabling the model to learn proximity between homologous genes across species. For noncoding or species-specific genes without homologs, we annotated gene types explicitly to allow the model to distinguish species-specific expression patterns.This multi-layered strategy enabled the model to learn both conserved molecular programs and species-specific transcriptional signatures in a unified framework.

### Evaluation datasets

#### Heffel2024 (MERFISH, human hippocampus and cortex)

We used a spatial transcriptomics dataset profiling 29,691 tissue sections from the human hippocampus and prefrontal cortex across developmental time points using MERFISH^23^. This dataset is part of a multimodal resource that includes single-nucleus methyl-3C sequencing, designed to capture 3D chromatin conformation and gene expression during neurodevelopment. For our study, we used the MERFISH-derived gene expression data to evaluate zero-shot representation performance in developing human brain tissues. Raw and processed data are publicly available from GEO under accession GSE21395.

#### Fang2022 (MERFISH, human cortex)

We used a MERFISH-based spatial transcriptomics dataset profiling the middle and superior temporal gyrus of the human cerebral cortex^24^. The dataset includes 101,201 cells and 3,293 genes across 15 tissue sections from four healthy donors. It provides a high-resolution atlas of transcriptionally distinct cell populations and spatial architectures, including neuron-glia interaction patterns and conserved laminar organization. This dataset was used to evaluate zero-shot representation performance in complex human cortical regions. Raw and processed data are publicly available via Dryad (https://datadryad.org/stash/dataset/doi:10.5061/dryad.x3ffbg7mw) and SODB.

#### Fang2022 (MERFISH, mouse cortex)

We also included the corresponding mouse dataset from the same study^24^, which profiles 26,072 cells and 237 genes across two tissue sections from two healthy donors, covering auditory, temporal association, and visual cortical areas. This dataset was used to assess cross-species generalization under consistent MERFISH protocols, enabling comparison of laminar and intercellular organization between mouse and human cortices. Raw and processed data are publicly available via Dryad (https://datadryad.org/stash/dataset/doi:10.5061/dryad.x3ffbg7mw) and SODB.

#### Allen2022-Aging (MERFISH, mouse cortex and striatum)

We used a MERFISH-based spatial transcriptomics dataset characterizing molecular and cellular aging in the mouse frontal cortex and striatum^25^. The dataset includes 378,918 cells and 374 genes across 31 coronal sections from 12 donors spanning the mouse lifespan. It provides detailed spatial expression profiles and captures age-associated transcriptomic alterations across major neural cell types. This dataset was used to evaluate zero-shot representation performance across physiological aging conditions. Raw and processed data are publicly available via CellxGene (https://cellxgene.cziscience.com/collections/31937775-0602-4e52-a799-b6acdd2bac2e) and SODB.

#### Allen2022-LPS (MERFISH, mouse cortex and striatum)

We also used a companion dataset from the same study^25^ profiling inflammation-induced changes in the mouse brain following systemic lipopolysaccharide (LPS) challenge. This dataset comprises 345,934 cells and 374 genes across 18 coronal sections from 8 donors. It captures distinct spatial transcriptomic signatures of glial and immune cell activation and was used to assess the model’s generalization to acute neuroinflammatory perturbations. Raw and processed data are publicly available via CellxGene (https://cellxgene.cziscience.com/collections/31937775-0602-4e52-a799-b6acdd2bac2e) and SODB.

#### Ma2024-Aging (Stereo-seq, mouse brain)

We used the brain subset of a Stereo-seq-based spatial transcriptomics dataset profiling aging-associated transcriptomic changes across multiple mouse organs^26^. The dataset provides ultra-high-resolution spatial expression maps and identifies senescence-sensitive spots (SSSs) with local accumulation of immunoglobulin-expressing cells and increased structural entropy. For our study, only the brain sections were used to evaluate the model’s generalization across aging phenotypes under high-resolution spatial profiling. Raw and processed data are available via the Cell publication and associated resources.

### BrainBeacon architecture and pretraining

BrainBeacon is a Transformer-based foundation model designed for cross-species and cross-platform representation learning on spatial transcriptomic data. To capture both molecular features and spatial context, BrainBeacon adopts a hierarchical two-stage architecture that models intra-cellular gene expression and inter-cellular spatial relationships. In the first stage, each cell is encoded as a fixed-length sequence of gene tokens, which are processed by a gene-level Transformer to learn molecular representations. In the second stage, cells within the same tissue slice are treated as cell tokens and jointly modeled using a separate Transformer to capture spatial organization and tissue structure.

This two-stage design enables BrainBeacon to generalize across diverse technologies and species while preserving biologically meaningful representations. In the following sections, we describe the tokenization strategy, intra- and inter-cell Transformer modules, and the training procedure in detail.

#### Gene tokenization strategy

To represent each cell as a structured input for intra-cellular modeling, BrainBeacon encodes its transcriptomic profile into a fixed-length token sequence that incorporates both biological prior knowledge and contextual metadata. Each sequence consists of three auxiliary tokens: species, platform, and spatial cell density. In Addition, each sequence is followed by the top-*K* most highly expressed genes ranked by gene expression value descendingly.To reduce platform-specific biases and improve robustness to expression sparsity, we normalize each gene’s expression using platform-specific non-zero means:

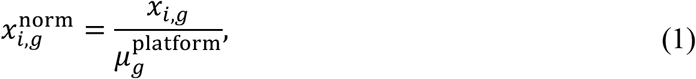

where 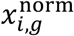 denotes the raw count of gene *g* in cell *i*, and 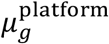 is the mean non-zero expression of gene *g* across all cells from the same platform. The top *K =* 997genes with the highest normalized expression are retained to ensure a fixed input length of *T* = 1000, including the 3 auxiliary tokens. If a cell contains fewer than 997 non-zero genes, special padding tokens are added to maintain sequence length.

Each token *t* ϵ {1, …, 1000} is mapped to a unified embedding vector by summing relevant components:

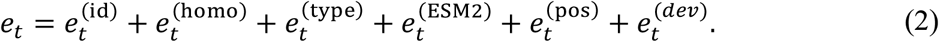

Here, 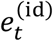 is the gene ID embedding, assigned from a shared vocabulary of 92076 Ensemble gene identifiers across species. 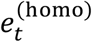 represents homology group embeddings, derived from BioMart-based orthology graphs that define minimal cross-species connect components. 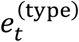 denotes RNA type embeddings reflecting gene category annotations (e.g.,protein-coding, lncRNA). 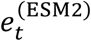 denotes the fixed protein-level embedding for protein-coding genes, derived from pretrained ESM2 model^27^ and subsequently projected through a multilayer perceptron (MLP) to align with the dimensionality of 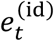. Finally, 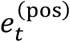 is a sinusoidal positional encoding indicating each token’s order within the sequence, and 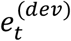 represents the niche-aware deviation of each gene’s expression, computed as the difference between a cell’s expression and the average expression of its spatial neighbors within a fixed radius, and discretized into quantile bins.

The auxiliary tokens for species, platform, and cell density are appended to the beginning of the input sequence and encoded through dedicated embedding lookup tables. The species token informs the model of the organism of origin (e.g., human, mouse, macaque, marmoset); the platform token encodes the measurement technology (e.g., MERFISH, Slide-seqV2, Xenium); and the cell density token reflects local spatial context, computed by counting neighboring cells within a fixed radius and discretized into ordinal bins.

All token embeddings are stacked to form the input matrix for a given cell:

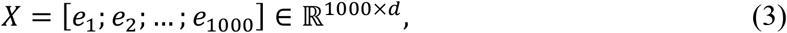

where *d* is the embedding dimension (default 1024). This matrix is then fed into the gene-level Transformer encoder to learn context-aware molecular representations.

This tokenization strategy allows BrainBeacon to encode heterogeneous spatial transcriptomic data from diverse species and platforms, while preserving spatial context, functional gene properties, and evolutionary structure.

#### Intra-cellular modeling with Transformers

Following tokenization, each cell is represented as a fixed-length sequence of *T* = 1000 tokens, including auxiliary and gene tokens. These sequences are passed through a multi-layer Transformer^28^ encoder, which models context-dependent gene relationships using stacked self-attention blocks.

The input sequence is embedded as a matrix*X* ϵ ℝ^*T×d*^, and contextualized representations are obtained by applying stacked Transformer layers:

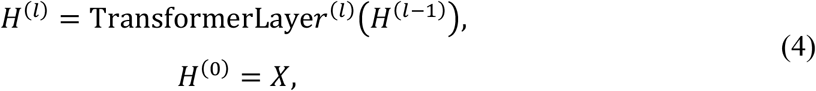

where each Transformer layer contains multi-head scaled dot-product attention and position-wise feedforward modules. The self-attention mechanism in each head operates as:

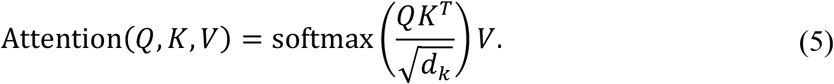

with *Q, K*, 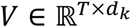 obtained from learned linear projections of the input embeddings.

To train the encoder, we adopt a masked gene prediction (MGP) objective, analogous to masked language modeling. A subset ℳ ⊂ {4, …, D}of gene token positions is randomly selected for masking (auxiliary tokens at positions 1-3 are excluded), and the model is tasked with recovering their identities from context:

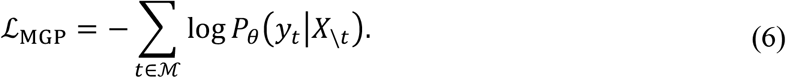

Here, *y*_t_ is the ground-truth gene ID at position *t, X*_\t_ denotes the input sequence with token *t* masked, and *P*_θ_ is the model-predicted probability over the gene vocabulary using a *softmax* classifier. This objective enables the encoder to learn gene–gene dependencies within each cell, producing context-aware representations for downstream spatial analysis.

#### Inter-cell input construction

To model spatial relationships among cells, BrainBeacon constructs cell-level token sequences for each tissue slice and applies a Transformer encoder to capture inter-cellular dependencies. Each cell token is derived from the gene-level contextual representations obtained via the intra-cell Transformer encoder, aggregated into a single vector through an expression-guided weighted pooling mechanism. Specifically, the aggregation includes all valid tokens except for the platform, species, and padding tokens, and retains the cell density token.

We denote the final output of the gene-level Transformer for token *t* as 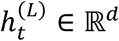, where *L* is the total number of layers. Formally, for a given cell *i*, the input cell embedding 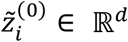 is computed as:

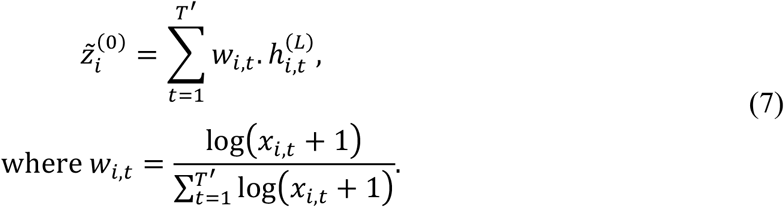

Here *x*_*i,t*_ denotes the platform-normalized expression value for gene token *t* in cell *i*, as defined in equation (1), and *T*′ is the number of included tokens in the sequence (i.e., excluding species, platform, and padding tokens only).

To incorporate spatial context, the 2D coordinate of cell *i* is encoded using a sine-cosine positional embedding^16^ *p*_*i*_ ϵ ℝ^*d*^. The final input embedding of cell *i* s defined as the sum:

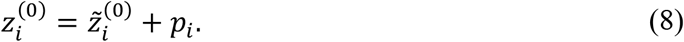

All cell embeddings 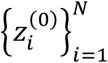 are stacked row-wise to form the input matrix *Z*^(0)^ ϵ ℝ^*N×d*^,where *N* is the number of cells within the brain slice. The matrix *Z*^(0)^ is then fed into the inter-cell Transformer encoder to model spatial dependencies.

#### Inter-cellular modeling and spatial expression reconstruction

To capture spatial dependencies and enable expression-level inference under missing or perturbed conditions, BrainBeacon integrates a masked cell modeling strategy with a variational autoencoding framework. During training, a random subset of cells is selected as reconstruction targets (masked cells), and the model is trained to recover their full gene expression profiles.

For each masked cell *i*, a variational Transformer encoder produces the parameters of a latent Gaussian distribution 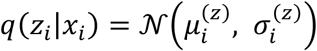, where the encoder integrates expression and spatial context information from neighboring cells to compute the latent representation. The latent representation *z*_i_ ϵ ℝ^d^ is sampled using the reparameterization trick. The latent vector is then decoded via a two-layer multilayer perceptron (MLP) to output gene-wise parameters (*μ*_*ig*_, *θ*_*ig*_) ϵ ℝ^*G*^ × ℝ^*G*^, representing the predicted mean and inverse dispersion parameters of the negative binomial (NB) distribution for gene *g* in cell *i*.

The model is optimized to reconstruct the expression of masked cells using an NB-based likelihood loss:

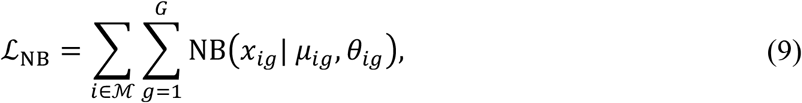

where ℳ denotes the set of masked cells and *x*_*ig*_ is the observed count of gene *g* in cell *i*. A Kullback–Leibler (KL) divergence term is used to regularize the approximate posterior against a standard normal prior:

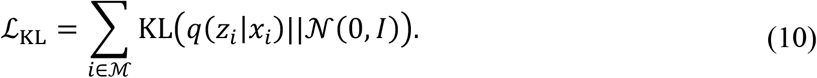

The total training objective of this stage is given by:

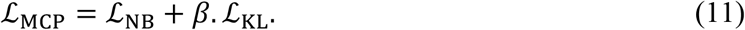

where *β* is a weighting coefficient.

The model is trained to reconstruct masked cells using Transformer-learned attention across the slice, enabling context-aware and uncertainty-aware inference in partially observed spatial environments.

#### Pretraining implementation

BrainBeacon adopts a two-stage training strategy, including intra-cell pre-training by masked gene prediction and inter-cell pre-training by spatial celluar reconstruction. In the first stage, 15% of gene tokens are randomly masked from a vocabulary of 92076, with supervision provided by 46502 homology-based connected components. Each token is embedded into a 1024-dimensional space and processed by a 16-layer Transformer encoder, with each layer featuring 1024 hidden units and 16 attention heads. To address distributional biases across species and sequencing platforms, samples are drawn with probabilities inversely proportional to their joint occurrence frequency, thereby promoting balanced representation. The model is optimized using AdamW with a learning rate of 0.00001 and an effective batch size of 1024 over 6 epochs.

In the second stage, the pre-trained cell embeddings *Z*^(0)^ are used to reconstruct observed gene expression profiles. A 4-layer Transformer encoder with 1024 hidden units and 8 attention heads is employed as the backbone, followed by a 2-layer multilayer perceptron (MLP) to project the outputs for loss computation. This stage is trained for 100 epochs using AdamW with a learning rate of 0.0002.

All training procedures are conducted on a computing cluster comprising 64 NVIDIA A100 GPUs across 8 nodes, each equipped with 120 CPU cores and 2 TB of RAM.

### Downstream tasks

#### Zero-shot embedding and annotation

To evaluate the generalization capability of BrainBeacon representations, we perform downstream tasks in a zero-shot setting, without any fine-tuning on the target datasets. We utilize the gene-level embeddings 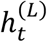 produced by the intra-cell Transformer and the cell level embeddings *z*_*i*_ output by the inter-cell Transformer as universal representations.

For cell-level tasks, such as clustering or annotation, we directly use the cell embeddings *z*_*i*_ ϵ ℝ^*d*^ computed from the target slice. These embeddings capture both intrinsic transcriptional identity and spatial context, and are expected to generalize across species and platforms.

For gene-level tasks, such as marker discovery or cross-slice comparison, the token-wise embeddings 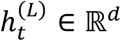 provide context-aware representations of gene activity, incorporating both expression value and gene identity information.

In label transfer applications, we fine-tune the model on labeled snRNA-seq data and apply it to target ST slices. Cell embeddings *z*_*i*_ ϵ ℝ^*d*^ are projected through the trained classification head to produce logits, which are refined using anti-collapse strategies to yield final cell type predictions.

#### Label transfer across platform and species

We implemented a cross-platform label transfer module built on the BrainBeacon architecture to propagate cell-type annotations from well-annotated snRNA-seq references to unlabeled spatial transcriptomics (ST) datasets. The model is first fine-tuned on the reference dataset using a classification head trained atop the pre-trained BrainBeacon encoder. During fine-tuning, we freeze early embedding layers to prevent overfitting and apply Focal Loss to address class imbalance in the reference.

For inference on ST datasets, we adopt a temperature-scaled softmax over the model’s output logits to generate class probabilities. To mitigate the risk of prediction collapse caused by overrepresented classes, we integrate a universal anti-collapse module. This module applies entropy regularization to increase output diversity, confidence sharpening to enhance high-certainty predictions, and adaptive suppression to down-weight dominant classes in the output distribution. These operations are guided by a dynamically selected strategy profile (e.g., “conservative”, “aggressive”, “ultra-balanced”), chosen based on class imbalance metrics such as the number of classes and imbalance ratio observed in the reference set.

To further improve generalization, we monitor prediction uncertainty and distribution skew by computing the average *softmax* entropy and the normalized class entropy on the target ST data. These metrics are used to adjust inference parameters such as *softmax* temperature and suppression strength, enabling the model to adaptively calibrate predictions under varying class priors and improve robustness to domain shifts.

#### In silico spatial perturbation of cellular and microenvironmental contexts

To evaluate BrainBeacon’s capacity to model context-aware cellular state changes, we implemented two types of in silico perturbations: cell-centric (*cell2niche*) and microenvironmental (*niche2cell*). These perturbations simulate biological variation at both intracellular and niche levels, enabling systematic assessment of the model’s response to virtual interventions.

In the *cell2niche* setting, we directly perturb a subset of target cells by activating, suppressing, or masking selected genes, and monitor the resulting changes in cell embeddings and reconstructed gene expression. This simulates molecular-level interventions such as genetic mutations or therapeutic modulation. Perturbations are applied by modifying either the normalized expression profiles or the ranked gene tokens of the selected cells prior to model inference. The updated representations capture both the direct impact on perturbed cells and the indirect influence on their neighbors through spatial attention.

In the *niche2cell* setting, we manipulate the local microenvironment by adding, removing, or replacing neighboring cells, while keeping the target cell unchanged. This enables the investigation of cell–niche interactions, such as immune cell infiltration or the effect of neighboring senescent cells. To simulate such contexts, we construct modified spatial slices by introducing donor cells into the vicinity of the target cell, preserving spatial coordinates while altering the surrounding contextual signals.

## Data availability

The datasets used and analyzed in this study will be made publicly available upon publication.

## Code availability

The BrainBeacon code and pretrained models will be made publicly available upon publication. Pre-release access may be granted upon reasonable request.

## Acknowledgements

This work was supported by National Natural Science Foundation of China (Nos. T2341007, T2350003, 12131020, 42450084, 42450135, 12326614, 12426310, 62002329, 82394432 and 92249302); National Key R&D Program of China (2022YFA1004800, 2025YFF1207900); Science and Technology Commission of Shanghai Municipality (23JS1401300); Shanghai Leading Talent Program of Eastern Talent Plan to W.W.; Zhejiang Province Vanguard Goose-Leading Initiative (2025C01114); Hangzhou Institute for advanced study of UCAS (2024HIAS-P004); Major Science and Technology R&D Project of the Science and Technology Department of Jiangxi Province [20213AAG01013]; JST (Japan Science and Technology Agency) Moonshot R&D (No. JPMJMS2021); AMED under Grant Number JP23dm0307009; Institute of AI and Beyond of the University of Tokyo; International Research Center for Neurointelligence (WPI-IRCN) at the University of Tokyo Institutes for Advanced Study (UTIAS); JSPS KAKENHI Grant Number JP20H05921;Henan Province Natural Science Foundation (242300421401); Open Research Fund of State Key Laboratory of Digital Medical Engineering (2024-K07); National Natural Science Foundation of China (Grant No. 82394432 and 92249302) and Shanghai Municipal Science and Technology Major Project (Grant No. 2023SHZDZX02).

## Ethics declarations

Not applicable

## Competing interests

The authors declare no competing interests.

## Notes

### Competing Interest Statement

The authors have declared no competing interest.

